# PRELIMINARY IN VITRO EVALUATION OF NARINGIN AND QUERCETIN FOR POTENTIAL BIOENHANCEMENT OF ANDROGRAPHOLIDE USING PLASMA PROTEIN BINDING AND HEPATIC MICROSOMAL STABILITY ASSAYS

**DOI:** 10.64898/2026.08.06.743340

**Authors:** Pragnya Pisipati, Tanmay Paranjpe, Shriya Natu, Aamir Khan, Vasudha Salgotra

**Author notes:** These authors contributed equally to the work (Pragnya Pisipati, Tanmay Paranjpe, Shriya Natu, Aamir Khan).

## Abstract

Several potentially potent anticancer drugs have been identified by in vitro evaluation, such as Andrographolide. These compounds show strong anticancer activity in vitro, but struggle to reach effective concentrations in the bloodstream when taken orally because they dissolve poorly in water or break down rapidly in the body. Bioenhancers, which are compounds that have potential to improve drug stability in the body, offer an alternative solution to overcome this limitation. Naringin and Quercetin have been identified as candidate bioenhancers, and have been hypothesized to potentially slow rapid first pass metabolism of poorly bioavailable drugs. Our work focuses on testing Naringin and Quercetin because they are flavonoids with therapeutic potential, due to their anti-inflammatory and antioxidant properties. Data from the hepatic microsomal assays performed on Naringin and Quercetin suggest moderate to proficient periods of stability in the body, with Naringin having 91.86% remaining, while Quercetin had 74.84% remaining. When administered alongside Andrographolide, a drug known to rapidly degrade in the body, Naringin raised its metabolic stability from 38.93% to 80.77% and on the other hand, Quercetin raised Andrographolide metabolic stability from 38.93% to 86.70%. In addition, plasma protein binding assays show the percentage of compounds available at the target site where Naringin was observed to be 49.32% bound and Quercetin found to be 50.14% bound, implying 50.68% of Naringin, and 49.86% of Quercetin available at the target site, respectively. This preliminary study explores whether Quercetin and Naringin could act as bioenhancers by remaining stable and available in plasma and by slowing the metabolism of poorly bioavailable drugs such as Andrographolide.

## Introduction

With the introduction of many new therapeutic drugs that have the ability to target Poly-glycoproteins[1], utilizing these drug molecules has become an important part of pharmaceutical and pharmacokinetic research, which are the studies of the effects of drugs on the body and the movement of drugs around the body[2][3][4]. Poly-glycoproteins are a vital efflux protein of the cell that monitors the foreign substances that exit the cell[5][6]. They are responsible for different kinds of immune responses, cellular signaling, and the regulation of drug transport across several kinds of membranes [6]. The inhibition of Poly-glycoproteins has become a central topic in the pharmaceutical community, due to its ability and potential to overcome MDR, also known as multidrug resistance[7]. This is where cancer cells start to become impervious to many current chemotherapeutic treatments[8][9][10]. This study aims to investigate the potential of candidate bioenhancers quercetin and naringin, 2 bioactive molecules commonly found in fruits and vegetables[11][12]. The aim here is not to evaluate their P-gp inhibition potential, but to investigate their likely availability in the body and their potential to improve the metabolic stability of low-bioavailability drugs in the face of liver microsomes. We seek to work towards improving the state of current drug therapies, thereby contributing to future studies on preventing or reversing multidrug resistance (MDR) in cells.

Our study uses methods such as plasma-protein binding [13][14][15] to determine the binding ability of quercetin and naringin to plasma proteins and find their percent bound, indicating their possible availability at the cell target site after making their way through the bloodstream. We also conduct hepatic microsomal stability assays[16][17], which allow us to determine how long certain compounds last in the bloodstream. It helps us determine the metabolic stability of the compound and how long it would be until the compound is metabolized by the liver.

Poly-glycoproteins, or p-gps, are membrane proteins that work to remove and eject foreign molecules from the cell [5][6]. They are efflux pumps dependent on ATP[18], the source of energy at a cellular level[19]. P-gp controls the ability to absorb, distribute, and remove many medical compounds[5], many of which are being used to treat cancer. This removal of the chemotherapeutic agents via p-gp’s MDR creates a huge barrier for the potential recovery and treatment of patients[20].

**Figure 1:**
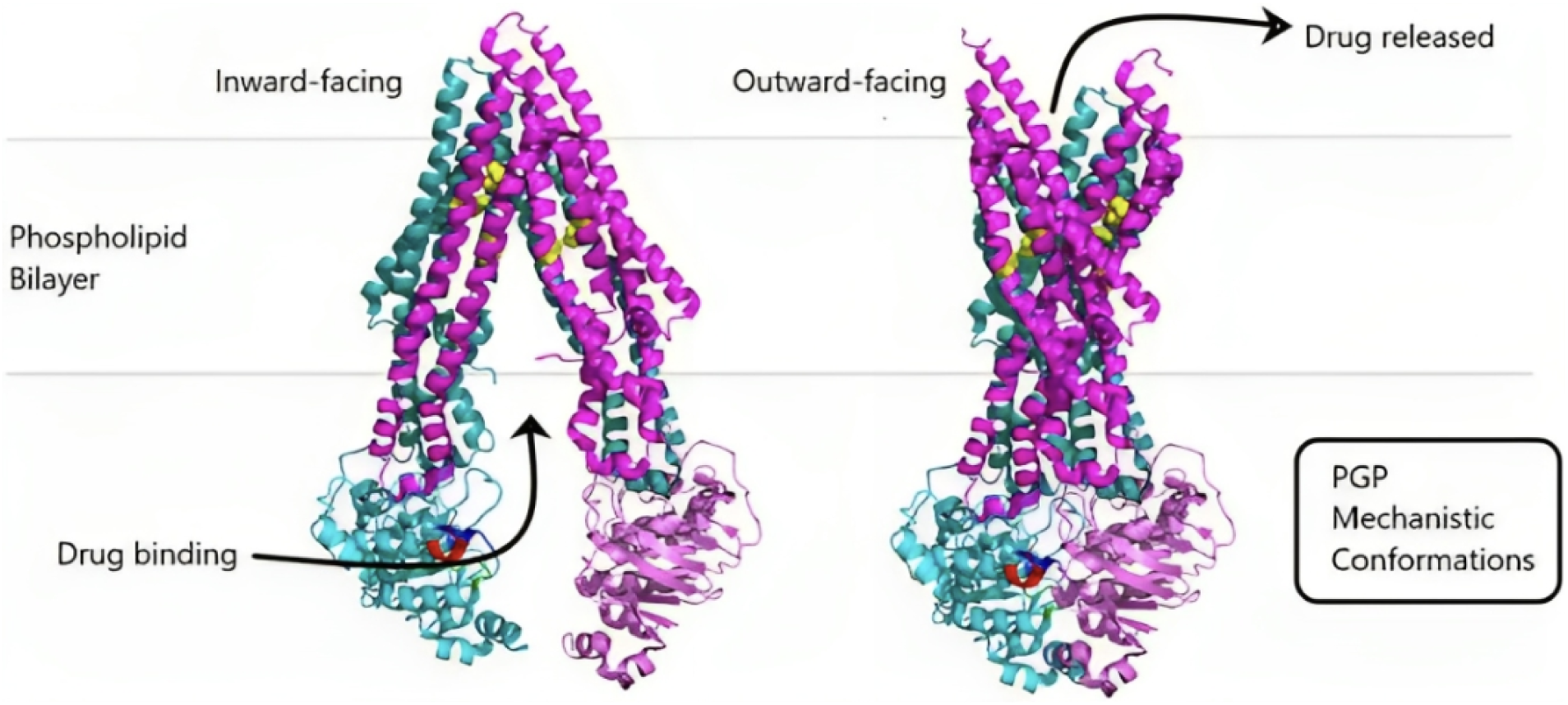
Depiction of Poly-glycoprotein and its mechanisms.

With cancer research becoming the centralized focus of pharmaceutical studies, many have focused their attention on being able to stop the abnormal growth of cells. Some have chosen to focus on the absorbance properties of the molecules, or also the bioavailability of the drug molecules. Previous research identifies that due to the efflux pump mechanism of poly-glycoproteins in cells, many chemotherapeutic agents are rejected from the cell. To resolve this, a search for a poly-glycoprotein inhibitor, or just a P-gp inhibitor. Many inhibitors have already begun to be developed and processed in order to increase the absorption of chemotherapeutic agents. In the past, researchers have already begun the search for a new inhibitor. In 2009, the Canadian Journal of Pharmacy came to the conclusion that flavonoids were a viable and possible P-GP inhibitor and could be used to allow anti-cancer drugs to become more bioavailable[21]. Even earlier, in 2002, researchers were able to find that Quinazolinones were a possible P-GP inhibitor, and they could be used to prevent MDR, or multidrug resistance[22]. Many researchers have been able to find several P-GP inhibitors, and this study has been an area of focus for many years, as it is vital to finding a much more effective chemotherapeutic agent.

Cancer, in the 21st century, has become one of the largest and deadliest diseases to plague the planet. It is essentially a tumor, or a group of cells, that randomly grows throughout the body. Its random and erratic movements make it hard to predict the disease. However, it is still possible to slow down or prevent the cancerous tumour. To possibly stop the growth of cells, we have developed chemotherapeutic agents, which kill off the abnormal cells. There are many different types of chemotherapeutic agents already being used, such as cyclophosphamide or doxorubicin[23][24]. As discussed previously, poly-glycoproteins are efflux pump proteins. They are responsible for removing foreign substances that do not belong in the cell. This can result in many anticancer agents that are used today being ineffective due to the aforementioned efflux function, which ejects the agent from the cell[25]. This causes a lack of treatment for the patient, which could severely threaten their life. To resolve this, we need a molecule that is able to stop the poly-glycoprotein from ejecting the chemotherapeutic agent. This would be called a P-GP inhibitor. To find one, we have to search for a candidate bioenhancer molecule that can stay available and stable in the body for extended periods of time and act in the desired mechanism to encourage drug stability in the body.

However, this is a widely studied problem within the pharmaceutical community. Different institutions have found different ways to be able to inhibit poly-glycoproteins. For example, researchers at the European Journal of Medicinal Chemistry have found that it is possible to use natural alkaloids as poly-glycoprotein inhibitors[26]. They have found that alkaloids, which are found in plants and different kinds of vegetation, have strong compatibility with chemotherapeutic agents, such as paclitaxel[27][28]. However, alkaloids fail to be able to strongly bind with proteins, allowing a small window for chemotherapeutic drugs to pass through[29]. On top of that, alkaloids are not very well tested in the scientific field. They are a new topic covered, and we have just recently started testing alkaloids for a possible P-GP inhibitor. However, there are other natural products that have shown potential for P-GP inhibition. For example, The Advanced Pharmacognosy Research Laboratory in India has discovered that a wide range of natural substances are viable P-GP inhibitors, such as terpenoids and saponins[30].

The molecule naringin was more recently discovered to have the potential to be a bioenhancer, with observed similarities to quercetin. Studies on naringin have shown that naringin is able to modulate the activities of drug-metabolizing transporters, like Poly-glycoproteins[31]. In studies on animals, naringin has shown enhancement in the absorption of some drugs and an increase in bioavailability as well[32]. This is through inhibiting the protein, which, in turn, stops the MDR from ejecting molecules from the cell[33].

**Figure 2:**
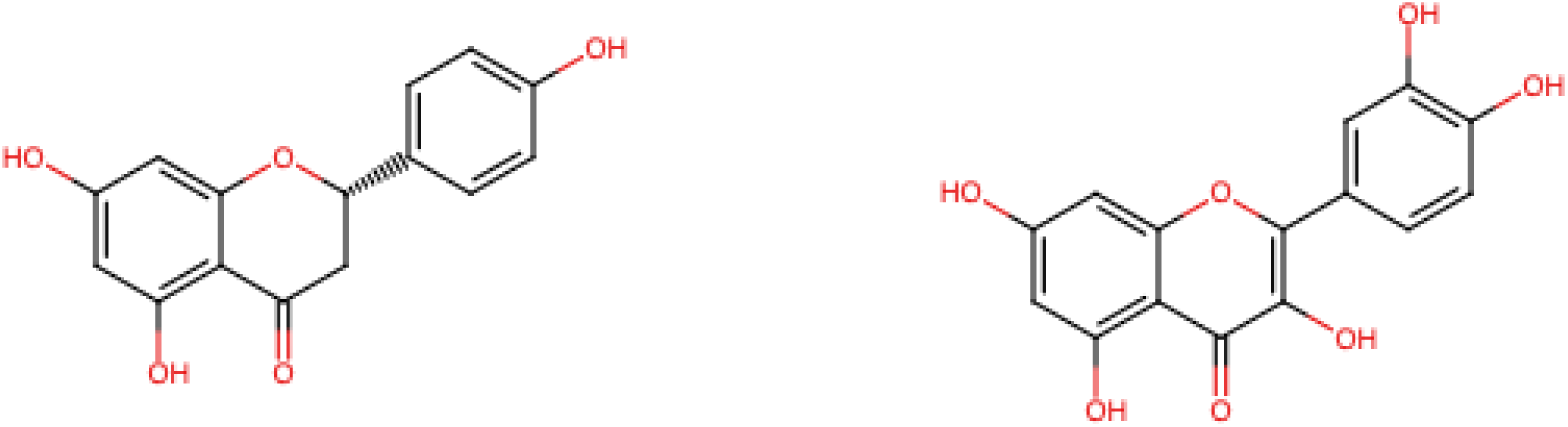
The chemical structure of naringin and quercetin, with naringin on the left and quercetin on the right, made using RCSB PDB Chemistry Sketch Tool

Quercetin, on the other hand, has been a studied and well-known molecule, capable of being able to interact with different proteins and enzymes[34]. Studies have been able to show that quercetin is able to inhibit P-gps, allowing the concentration of administered chemotherapeutic cells to be increased[35]. This allows a more effective dosage of drugs. Quercetin has been shown to have properties that increase the cytotoxicity of chemotherapeutic drugs[36].

The concept of using natural compounds to boost drug bioavailability without altering the drug itself is not new. Naringin and quercetin have both been identified as part of this broader class of natural potential bioenhancers, alongside compounds like piperine, genistein, and glycyrrhizin[37]. Prior work has explored quercetin’s bioenhancing effect on compounds such as curcumin and rosuvastatin, while naringin has been studied for its ability to inhibit P-gp and breast cancer resistance protein (BCRP) to improve drug absorption[37]. However, much of this existing research has focused on quercetin and naringin as bioenhancers for pharmaceutical drugs or isolated cancer-related pathways, with little investigation into their combined use with other poorly bioavailable natural compounds, such as andrographolide. Additionally, direct comparative data on their plasma protein binding and hepatic metabolic stability, side by side, under matched experimental conditions, is limited. Our study aims to fill this gap by directly evaluating naringin and quercetin’s plasma protein binding and hepatic microsomal stability using the same assay conditions, and testing their possible bioenhancing effect on andrographolide specifically, providing new data on how these compounds perform as candidate bioenhancers.

Andrographolide, the primary bioactive diterpenoid found in *Andrographis paniculata*, has drawn attention for its anti-inflammatory, antiviral, and anticancer properties[38]. Despite this therapeutic potential, andrographolide has notably poor oral bioavailability, largely due to rapid first-pass metabolism and active efflux by P-gp, with reported absolute bioavailability as low as 2.67%[38]. This makes andrographolide a strong candidate compound for testing the bioenhancing potential of our select compounds; its low natural bioavailability offers an opportunity for improvement through coadministration of potential bioenhancers. Naringin and quercetin, given their proposed possible bioenhancing activity, may help slow clearance of andrographolide and increase the proportion of the compound that remains available in the body over time. Testing naringin and quercetin alongside andrographolide therefore serves useful for evaluating whether these flavonoids can applicably function as bioenhancers, extending their relevance to other poorly bioavailable natural compounds.

Naringin and quercetin’s ability to overcome multidrug resistance could contribute to improving the bioavailability of certain chemotherapeutic drugs. Compared to previous synthetic bioenhancers that have been shown to exhibit significant toxicity[39][30], naringin and quercetin offer a potentially safer alternative. However, their efficacy and safety are still being tested in clinical trials. Quercetin and naringin are widely available in common dietary sources, which allows them to be easy candidates for further studies.[40][41][42][43]. These two molecules possess great potential in their ability to control and target oxidative stress and inflammation[44]. This allows an enhanced role in their therapeutic potential in cancer cells, due to the large part that inflammation and oxidative stress[44] play a key role in the growth of cancer.

This paper will discuss the possibility of the usage of naringin and quercetin molecules to slow rapid drug clearance and their potential ability to act as bioenhancers.

Here, we aim not to use the molecules quercetin and naringin as poly-glycoprotein inhibitors, but to evaluate the viability of the compounds in the body and the extent of their ability to improve metabolic stability of drugs with low bioavailability. Our goal is to utilize the low-cost, naturally-sourced compounds to provide worldwide healthcare against cancer to all. This research could also be a stepping stone for many other researchers on this topic. It offers a prerogative for future research into the direct interactions of naringin and quercetin with P-GP. It could be used to possibly apply to other illnesses or scenarios, and not just limited to cancer. Instead, it could be used to treat and help increase the bioavailability of many drugs, supporting future development of bioenhancing strategies.

## Methodology

The naringin and quercetin used in this study were supplied by the Sigma-Aldrich chemical sciences company. The nanodrop spectrophotometer used for data analysis was supplied by Thermofisher.

### Plasma Protein Binding Assay

Approximately 3.5–4 mL of plasma was thawed prior to the experiment. Stock solutions (10 mg/mL) of naringin, quercetin, and ibuprofen were prepared by dissolving 10 mg of each compound in 1 mL of dimethyl sulfoxide (DMSO) with thorough mixing to ensure complete dissolution. Working solutions were subsequently prepared in 1× phosphate-buffered saline (PBS) to final concentrations of 2.5 mg/mL for naringin, 1.5 mg/mL for quercetin, and 1 mg/mL for ibuprofen, which served as the positive control. The naringin working solution was prepared by diluting 250 μL of the stock solution with 750 μL of PBS, the quercetin working solution by diluting 150 μL of stock with 850 μL of PBS, and the ibuprofen working solution by diluting 100 μL of stock with 900 μL of PBS.

Rapid Equilibrium Dialysis (RED) devices were cleaned with isopropyl alcohol or ethanol, rinsed thoroughly with deionized water, and allowed to dry completely before use. Clean inserts were placed into the RED device and labeled according to the test compound. Each insert received 500 μL of thawed plasma in the plasma (red) chamber and 500 μL of the corresponding working solution in the buffer (white) chamber. Four experimental conditions were prepared: naringin, quercetin, ibuprofen (positive control), and a negative control consisting of 1× PBS without test compound. The RED device was sealed with a plate sealer and incubated at 37°C in a shaking incubator for 3 hours to allow equilibrium dialysis.

Following incubation, separate microcentrifuge tubes were prepared for the plasma and buffer samples corresponding to each experimental condition. A 300 μL aliquot was collected from both the plasma and buffer compartments of each insert and transferred to the appropriate labeled tubes. Protein precipitation was performed by adding 600 μL of methanol to each sample, followed by thorough mixing. The samples were centrifuged at 10,000 rpm for 10 min, after which the supernatants were carefully transferred into clean tubes labeled as plasma or buffer supernatants. The collected supernatants were subsequently analyzed using a NanoDrop spectrophotometer.

### Hepatic Microsomal Stability Assay

Hepatic microsomes were first thawed on ice prior to use. A 1 mM stock solution of NADPH in DMSO was prepared by weighing 0.744 mg of NADPH and dissolving it in 1,000 µL (1 mL) of DMSO. A 100 µM hydrocortisone solution was similarly prepared by dissolving 36.25 mg of hydrocortisone in 1 mL of DMSO. Naringin and quercetin solutions were each prepared at a concentration of 100 µM: 5.81 mg of naringin was dissolved in 1 mL of DMSO, and 3.02 mg of quercetin was dissolved in 1 mL of DMSO; each stock was then further diluted using 10 µL of stock solution combined with 990 µL of buffer. Andrographolide was prepared in the same manner, with 3.5 mg dissolved in 1 mL of DMSO and further diluted using 10 µL of stock solution combined with 990 µL of buffer, to yield a 100 µM working solution. A 100 mM MgCl₂ solution was prepared using anhydrous MgCl₂, with 9.521 g dissolved and brought to a final volume of 1 mL with deionized water; from this stock, 30 µL was combined with 970 µL of deionized water to prepare the working solution. Finally, a 5 mg/mL microsome stock was prepared by adding 50 µL of microsomes (20 mg/mL) to 150 µL of buffer, for a total volume of 200 µL.

Reaction mixtures were prepared for seven experimental conditions: a control containing no inhibitor, a positive control containing ketoconazole, naringin alone, quercetin alone, andrographolide alone, a combined naringin/andrographolide condition, and a combined quercetin/andrographolide condition. Each condition received 25 µL of microsomes (5 mg/mL), 7 µL of NADPH (10 mM), 12.5 µL of hydrocortisone (1 mM), and 3 µL of MgCl₂ (100 mM). The positive control additionally received 25 µL of ketoconazole (100 µM) and 177.5 µL of buffer. The quercetin-only condition received 25 µL of quercetin (100 µM) and 177.5 µL of buffer, while the naringin-only condition received 25 µL of naringin (100 µM) and 177.5 µL of buffer. The andrographolide-only condition received 25 µL of andrographolide (100 µM) and 177.5 µL of buffer. The combined naringin/andrographolide condition received 12.5 µL of naringin (100 µM) and 12.5 µL of andrographolide (100 µM), along with 177.5 µL of buffer, and the combined quercetin/andrographolide condition received 12.5 µL of quercetin (100 µM) and 12.5 µL of andrographolide (100 µM), along with 177.5 µL of buffer. The control condition, which received no inhibitor or test compound, was brought to volume with 202.5 µL of potassium phosphate buffer (pH 7.4). All conditions were prepared to a final total reaction volume of 250 µL.

Reaction mixtures were incubated at 37 degrees Celsius for a total of 90 minutes, with samples collected at three time points: 0, 60, and 90 minutes. At each time point, 3.5 µL of NADPH was added to the reaction to sustain enzymatic activity throughout the incubation period.

## Results and Discussion

### Plasma Protein Binding Assay

The viability of naringin and quercetin can be assessed using the binding percentage of molecules to proteins, specifically, via the Plasma protein-binding assay. This assay determines the binding potential of the molecules quercetin and naringin to plasma proteins and is used to assess what proportion of naringin and quercetin remains freely available rather than sequestered by plasma proteins, since only the unbound fraction of a compound is pharmacologically active and capable of interacting with P-gp or other target sites. A lower percent bound is therefore preferred to ensure that more compound is free to selectively bind to the target site or to poly-glycoproteins, minimizing off-target interactions. The data in Table 1 represents the percentage of each molecule bound to plasma proteins by the end of a two-hour incubation period. The fraction of molecule which was bound was calculated using the following equation: (C_total_-C_free_)/C_total_, where C_total_ is measured from the plasma chamber of the RED device and C_free_ is measured from the buffer chamber. This fraction bound was multiplied by 100 to get the percent bound.

**Table 1:** Depicts the results of three Plasma Protein Binding trials (i.e., the percentage of each molecule bound to plasma proteins) as well as the average percentage bound for each molecule and the corresponding standard deviations (mean±SD).

| Sample | Trial 1 | Trial 2 | Trial 3 | Average |
| --- | --- | --- | --- | --- |
| Naringin | 39.69% | 49.87% | 49.32% | 46.29%±5.73% |
| Quercetin | 30.00% | 50.00% | 50.14% | 43.38%±11.59% |
| Ibuprofen | 86.01% | 82.90% | 82.49% | 83.8%±1.92% |

Ibuprofen, used as the positive control due to its well-documented high plasma protein binding affinity, was 86.01%, 82.90%, and 82.49% bound across Trials 1, 2, and 3, respectively. This consistency across trials, with less than a 4% spread, indicates that the assay was performing reliably and that the RED device equilibration and NanoDrop quantification steps were functioning as expected across the full data collection window.

Naringin was more variable, with 39.69% bound in Trial 1 compared to a value of 49.87% and 49.32% in Trials 2 and 3. Quercetin had a similar pattern, with 30.00% bound in Trial 1 versus 50.00% and 50.14% in Trials 2 and 3. In both cases, Trial 1 differs from Trials 2 and 3 by roughly 10 percentage points, while Trials 2 and 3 are near in range. This suggests that Trial 1 could have possibly been affected by an inconsistency or environmental issue during experimentation. For example, an incomplete stabilization during the 2-hour incubation period, a pipetting inconsistency in the low volume dilution steps (as little as 10 µL of stock solution was used in several dilutions, which is difficult to pipette with high precision even with a calibrated micropipette), or a difference in how long the plasma had been thawed prior to use could have all caused the variation. Because Trials 2 and 3 are much more similar, we view them as the more reliable estimate of naringin and quercetin’s plasma protein binding behavior, while still reporting Trial 1 as part of the overall dataset. However, even with taking the lower Trial 1 values into account, naringin and quercetin were consistently far less protein-bound compared to ibuprofen across all three trials, which supports the conclusion that a significant amount of both compounds, meaning more than half, remains unbound and available to act on P-gp or other targets.

### Hepatic Microsomal Stability Assay

While the Plasma Protein Binding assay measures the extent to which Naringin and Quercetin will resist the processes of binding to proteins, understanding the stability of these molecules is a crucial factor in evaluating their properties as well. Stability of the molecules in the bloodstream is key to evaluating their metabolic stability, particularly in Phase 1 metabolism in the liver. This can be verified using the hepatic liver microsomal assay, which uses liver CYP450 enzymes to simulate Phase 1 oxidative metabolism in the liver and predict how quickly molecules are metabolized. The Hepatic Microsomal Stability assay evaluated how well naringin and quercetin, alone and in combination with andrographolide, withstand Phase I oxidative metabolism in the liver over a 90-minute period. A bioenhancer is expected to show a relatively high percentage remaining at the 90-minute mark on its own, and more importantly, to increase the percentage of a low-bioavailability compound remaining at the same time point. The data in the following tables (Tables 2, 3, and 4) represents the percentage of each molecule remaining unmetabolized at the end of each time point, with a table representing each trial. Furthermore, the data of each table is represented in a line graph to visualize the metabolization of each molecule over the 90-minute period.

**Table 2:** Depicts the results of Hepatic Microsomal Stability Assay Trial 1, with each percentage representing the percentage of the molecule remaining unmetabolized at the end of the specific time point.

| Sample | 0 minutes | 60 minutes | 90 minutes |
| --- | --- | --- | --- |
| Naringin | 100% | 80.60% | 62.70% |
| Quercetin | 100% | 99.10% | 62.20% |
| Andrographolide | 100% | 78.30% | 30.10% |
| Naringin +<br>Andrographolide | 100% | 79.20% | 59.50% |
| Quercetin +<br>Andrographolide | 100% | 82.90% | 55.30% |

**Table 3:** Depicts the results of Hepatic Microsomal Stability Assay Trial 2, with each percentage representing the percentage of the molecule remaining unmetabolized at the end of the specific time point.

| Sample | 0 minutes | 60 minutes | 90 minutes |
| --- | --- | --- | --- |
| Naringin | 100% | 95.05% | 91.58% |
| Quercetin | 100% | 72.86% | 55.71% |
| Andrographolide | 100% | 55.38% | 37.80% |
| Naringin + Andrographolide | 100% | 89.94% | 83.24% |
| Quercetin + Andrographolide | 100% | 85.71% | 84.69% |

**Table 4:** Depicts the results of Hepatic Microsomal Stability Assay Trial 3, with each percentage representing the percentage of the molecule remaining unmetabolized at the end of the specific time point.

| Sample | 0 minutes | 60 minutes | 90 minutes |
| --- | --- | --- | --- |
| Naringin | 100% | 96.51% | 91.86% |
| Quercetin | 100% | 95.55% | 74.83% |
| Andrographolide | 100% | 74.67% | 38.93% |
| Naringin + Andrographolide | 100% | 81.08% | 80.77% |
| Quercetin + Andrographolide | 100% | 92.26% | 86.70% |

**Table 5:** Depicts the average results of the Hepatic Microsomal Stability Assay trials, with each percentage representing the average percentage of the molecule remaining unmetabolized at the end of the specific time point across the 3 trials, along with the corresponding standard deviations (mean±SD).

| Sample | 0 minutes | 60 minutes | 90 minutes |
| --- | --- | --- | --- |
| Naringin | 100% | 90.72% ± 8.79% | 82.05% ± 16.76% |
| Quercetin | 100% | 89.17% ± 14.24% | 64.25% ± 9.72% |
| Andrographolide | 100% | 69.45% ± 12.32% | 35.61% ± 4.81% |
| Naringin + Andrographolide | 100% | 83.41% ± 5.74% | 74.50% ± 13.05% |
| Quercetin + Andrographolide | 100% | 86.95 ± 4.80% | 75.56% ± 17.58% |

Andrographolide, included specifically because of its previously documented poor bioavailability and rapid P-gp-mediated efflux, degraded rapidly across all three trials, falling to an average of 35.61% ± 4.81% remaining. This breaks down into 30.10% remaining in Trial 1, 37.80% in Trial 2, and 38.93% in Trial 3. This consistency, particularly between Trials 2 and 3, supports our selection of andrographolide as a suitable low-stability compound for testing the possible bioenhancing effects of naringin and quercetin, since its instability was reproducible rather than a one-off result.

Naringin alone showed a clear pattern across the three trials: 62.70% remaining at 90 minutes in Trial 1, but 91.58% and 91.86% remaining in Trials 2 and 3. The close agreement between Trials 2 and 3 (within 0.3 percentage points of each other) suggests that Trial 1 is the outlier here rather than Trials 2 and 3 being inconsistently high, and that naringin’s true stability profile is closer to the 92% range, with the compound largely resisting Phase I metabolism on its own. Quercetin alone followed a less consistent pattern, with 62.20%, 55.71%, and 74.83% remaining in Trials 1, 2, and 3, respectively. Unlike naringin, quercetin’s Trial 2 result does not clearly group with either Trial 1 or Trial 3, suggesting more inherent trial-to-trial variability in quercetin’s metabolic stability, or a data collection or handling issue specific to Trial 2’s quercetin condition, such as a delay in sampling at the 60-minute mark or a difference in how quickly the quercetin working solution was used after preparation given quercetin’s light and oxidation sensitivity in solution. For these reasons, the separate data for each trial is far more telling than the averages, though those are included as well to provide a more complete picture of the results.

The same pattern of Trial 1 being the outlier reappears in the conditions designed to mimic bioenhancer effects. When co-administered with andrographolide, naringin increased andrographolide’s percentage remaining from 30.10% (alone) to 59.50% in Trial 1, but from 37.80% to 83.24% in Trial 2 and from 38.93% to 80.77% in Trial 3. Quercetin produced a similar pattern, increasing andrographolide’s percentage remaining to 55.30% in Trial 1, but to 84.69% and 86.70% in Trials 2 and 3. In all three trials, the assisted andrographolide (co-administered with either naringin or quercetin) surpasses andrographolide alone by a large amount, meaning the direction of the bioenhancing effect held consistently regardless of trial. However, the extent of the observed effect was consistently smaller in Trial 1 than in Trials 2 or 3 across every sample in the assay, not just for one compound or one condition. These trends are exemplified by the shapes of the result graphs. This pattern across an entire trial points toward a trial level source of error like microsome batch activity, a delay between compound preparation and reaction start, or a temperature inconsistency during that specific incubation run, rather than a compound-specific issue with the tested substances.

### Interpreting Variability Across Trials

When observed alongside one another, the plasma protein binding assay and the hepatic microsomal stability assay show a similar pattern. Trial 1 invariably yielded lower stability and higher variability, while Trials 2 and 3 were correlated with one another across nearly every sample. This similarity may indicate that Trial 1, conducted earliest, could reflect an early stage methodological inconsistency. These inconsistencies may include a difference in reagent freshness, rather than biological variability in naringin and quercetin’s behavior. The conclusions that naringin and quercetin have low plasma protein binding compared to ibuprofen, and that both compounds significantly improve andrographolide’s metabolic stability when administered together, remained consistent across all three trials, regardless of variability. This finding suggests weighing Trials 2 and 3 more heavily as a more reliable representative of the compounds’ behavior, while continuing to include Trial 1 and utilizing it as a basis for improving consistency and replication.

### Limitations

Some limitations should be considered when interpreting these results:

1. Small number of trials: With only three trials per assay, we are not able to perform robust statistical significance testing to confirm that the differences observed between naringin, quercetin, and their controls are statistically meaningful rather than due to chance or experimental noise. Future work should aim for a minimum of five to six trials per condition to allow for meaningful statistical analysis and to better characterize discrepancies.
2. Potential spectral interference from NADPH and other reaction components: Because samples were quantified directly via NanoDrop spectrophotometry without a prior separation step, it is possible that NADPH or other molecules present in the reaction mixture (e.g., hydrocortisone, MgCl₂) absorb at wavelengths that overlap with those used to quantify naringin, quercetin, and andrographolide. NADPH in particular has a strong absorbance background, which could potentially slightly distort the analyte signal and introduce error into the reported percentages remaining at each time point. Because this was not directly tested in the current study, it remains a possible rather than confirmed source of error, but future work should consider troubleshooting this through using different data collection methods such as UV-Visible spectrophotometry and through troubleshooting via running negative controls.
3. Shaking speed control: Our incubations relied on a shaking incubator shared between researchers rather than a precision water bath, meaning small fluctuations in shaking speed during the 90-minute hepatic stability incubation or the 2-hour plasma protein binding incubation cannot be ruled out as a contributor to trial-to-trial variability.
4. Pipetting precision at low volumes: Several dilution steps in both assays required transferring volumes as small as 10 µL. At this scale, even a well-calibrated micropipette has a higher relative margin of error than it would at larger volumes.
5. Commercial microsome batch variability: Hepatic microsomes are a biological reagent, and enzymatic activity can vary between lots, storage conditions, and freeze-thaw history. We did not have the resources to verify baseline CYP450 activity of our microsome stock prior to each trial, which may partly explain the systematic difference observed between Trial 1 and Trials 2/3.

Despite these limitations, the consistency of the directional trends across all three trials (lower plasma protein binding for naringin and quercetin relative to ibuprofen, and a clear improving effect on andrographolide’s metabolic stability in every trial) provides encouraging preliminary support for their potential as bioenhancers that can stay viable in the body and improve the metabolic stability of low-bioavailability drugs such as andrographolide. The consistent and favorable findings warrant further investigation with larger sample sizes and more tightly controlled laboratory conditions. The outcomes also call for future research focusing on interactions of naringin and quercetin with P-gp directly in the cell, since their viability and bioenhancing potential in our assays thus far have been promising.

## Conclusion

This study provides preliminary evidence that naringin and quercetin may potentially function as candidate bioenhancers, with results that are consistent with, but do not demonstrate P-gp inhibition. However, in vivo testing is needed to confirm this. Across our plasma protein binding and hepatic microsomal stability assays, both flavonoids consistently remained less protein-bound than ibuprofen and substantially extended andrographolide’s stability against Phase I liver metabolism when co-administered. This is consistent with the hypothesis that they may reduce P-gp-mediated efflux and improve drug availability, although direct P-gp inhibition was not evaluated in this study. Given that naringin and quercetin are naturally sourced, readily available, and relatively low-toxicity compared to many established synthetic P-gp inhibitors, these findings point toward a promising, accessible route for improving the bioavailability of chemotherapeutic and other poorly absorbed compounds, with particular relevance to overcoming drug resistance in cancer treatment. Our next steps involve further controlled trials of hepatic microsomal stability and plasma protein binding to strengthen our findings. We would like to incorporate more enzymes in future hepatic stability assays to test for stability in Phase II liver metabolisation as well. Furthermore, we hope to work on cell permeability assays to test how well naringin and quercetin cross biological membranes and to evaluate their actual P-gp inhibition potential. In addition to further trials, future work also involves using UV-Visible spectrophotometry for more accurate low-concentration readings in order to rule out potential spectral interference from NADPH and other reaction components and confirm the accuracy of the reported concentrations. This confirmation can also be done by carrying out more thorough negative control trials, to compare the spectral signatures of NADPH and other reaction components by themselves to the spectral signatures of the components in the presence of naringin, quercetin, or andrographolide; this would allow us to distinguish our choice molecules’ readings from other components’ interference. Our work thus far has laid a groundwork for future studies to build upon, both in further validating naringin and quercetin’s bioenhancing potential and in extending this approach to other low-bioavailability compounds beyond andrographolide.

### Resource Management List

Our lead researcher and advisor was Dr. Vasudha Salgotra, and our collaborating researchers were Pragnya Pisipati, Tanmay Paranjpe, Aamir Khan, and Shriya Natu. We used the tools and resources of ASDRP, including the Nanodrop spectrophotometer and incubator, and molecules such as quercetin, naringin, ibuprofen, and andrographolide. We also used buffers and proteins from ASDRP’s stock. Since we were funded by ASDRP, we didn’t have a budget that we needed to adhere to on the project. We used Slack and Google Chat as our main sources of communication, and used the Google Scholar database to collect background information on our topic.

## Acknowledgements

We would like to thank Dr. Salgotra as our advisor who gave us immense support and direction. We would like to acknowledge Mrs. Gayatri Renganathan for providing us with the in silico data as the initial base which inspired our project and thank the rest of our team who are currently working on this project. We would also like to thank the rest of our team who previously worked on this project alongside us, including Chris Chen, Reva Ukkadam, and Tulika Sarkar. Last but not least we would like to thank ASDRP and the Olive Children Foundation for providing us the space and equipment to perform our research.

## Appendix

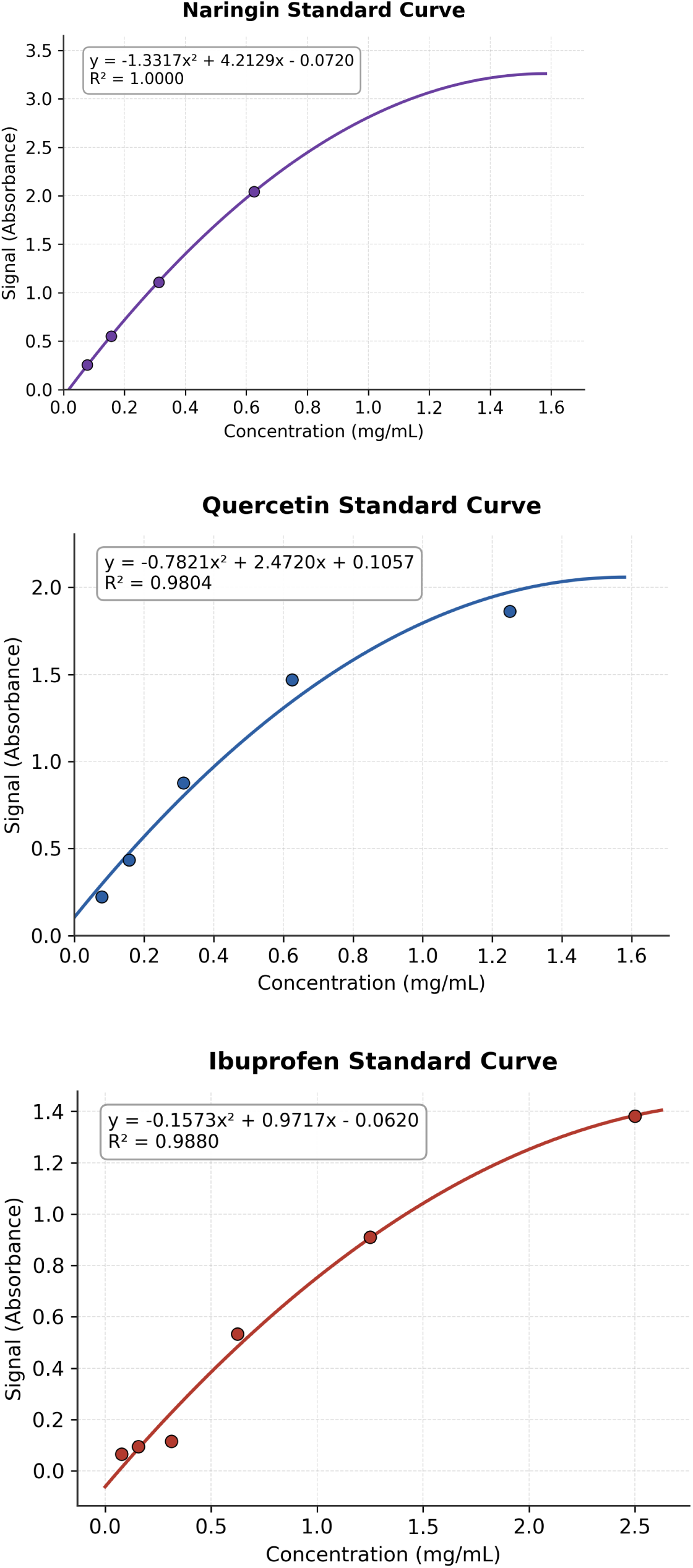

